# Mendelian randomization does not support serum calcium in prostate cancer risk

**DOI:** 10.1101/297721

**Authors:** James Yarmolinsky, Katie Berryman, Ryan Langdon, Carolina Bonilla, the PRACTICAL consortium, George Davey Smith, Richard M Martin, Sarah J Lewis

**Author notes:** Members from the Prostate Cancer Association Group to Investigate Cancer Associated Alterations in the Genome (PRACTICAL) consortium are provided in the Supplement notes. Information of the consortium can be found at http://practical.icr.ac.uk/. Corresponding author: Sarah J Lewis MRC Integrative Epidemiology Unit, School of Social and Community Medicine, University of Bristol, Oakfield House, Oakfield Grove, Bristol BS8 2BN, UK. GWAS: genome-wide association study HR: Hazard Ratio InSIDE: Instrument Strength Independent of Direct Effect IV: instrumental variable MR: Mendelian randomization OR: Odds Ratio PRACTICAL: Prostate Cancer Association Group to Investigate Cancer Associated Alterations in the Genome RR: Risk Ratio SNPs: single nucleotide polymorphisms WME: weighted median estimator.

## Abstract

**Background:** Observational studies suggest that dietary and serum calcium are risk factors for prostate cancer. However, such studies suffer from residual confounding (due to unmeasured or imprecisely measured confounders), undermining causal inference. Mendelian randomization uses randomly assigned (hence unconfounded and pre-disease onset) germline genetic variation to proxy for phenotypes and strengthen causal inference in observational studies.

**Objective:** We tested the hypothesis that serum calcium is associated with an increased risk of overall and advanced prostate cancer.

**Design:** A genetic instrument was constructed using 5 single nucleotide polymorphisms robustly associated with serum calcium in a genome-wide association study (N ≤ 61,079). This instrument was then used to test the effect of a 0.5 mg/dL increase (1 standard deviation, SD) in serum calcium on risk of prostate cancer in 72,729 men in the PRACTICAL (Prostate Cancer Association Group to Investigate Cancer Associated Alterations in the Genome) Consortium (44,825 cases, 27,904 controls) and risk of advanced prostate cancer in 33,498 men (6,263 cases, 27,235 controls).

**Results:** We found weak evidence for a protective effect of serum calcium on prostate cancer risk (odds ratio [OR] per 0.5 mg/dL increase in calcium: 0.83, 95% CI: 0.63-1.08; *P*=0.12). We did not find strong evidence for an effect of serum calcium on advanced prostate cancer (OR per 0.5 mg/dL increase in calcium: 0.98, 95% CI: 0.57-1.70; *P*=0.93).

**Conclusions:** Our Mendelian randomization analysis does not support the hypothesis that serum calcium increases risk of overall or advanced prostate cancer.

## Background

Prostate cancer is the most frequently diagnosed cancer among men globally and is a common cause of male cancer death [1]. Despite the considerable global burden attributed to prostate cancer, to date few risk factors (advanced age, ethnicity, family history of prostate cancer) have been identified and no modifiable risk factors have been established for this condition [2]. Nonetheless, global variation in prostate cancer mortality [3, 4] and findings from migration studies (i.e, the convergence toward local prostate cancer mortality rates among men who migrate from non-Western to Western populations) [5-7], provide support for a role of modifiable risk in prostate carcinogenesis.

Dietary calcium intake has been associated with an increased risk of prostate cancer in prospective epidemiological studies [8-10]. In a meta-analysis of fifteen prospective studies, high dietary calcium intake, as compared to low intake, was associated with an 18% (95% CI: 8-30%) increased prostate cancer risk [11]. Similarly, high calcium intake has been linked to an increased risk of advanced [12-14] and fatal prostate cancer [13], though findings have been inconsistent [15, 16]. Though serum calcium is normally tightly regulated in the body and does not fluctuate substantially across levels of dietary calcium intake [17, 18], Giovannucci proposed that higher dietary calcium may influence risk of prostate cancer by lowering circulating levels of 1,25(OH)2 vitamin D, a presumed tumour suppressor, [19-21] in order to achieve calcium homeostasis [22]. A more direct method of testing the hypothesis that calcium metabolism influences prostate carcinogenesis would be to examine the association of serum calcium levels with prostate cancer risk. However, studies examining the association of pre-diagnostic serum calcium levels with incident or fatal prostate cancer [23, 24], or post-diagnostic serum calcium with prostate cancer survival [25, 26], have generated conflicting results: some report positive associations of serum calcium with prostate cancer [23, 26, 27] whereas others have been compatible with a null effect [23-25, 28, 29]).

Establishing a causal role of elevated serum calcium in prostate carcinogenesis could have therapeutic implications for the prevention or treatment of prostate cancer. However, obtaining reliable estimates of causal effects from observational studies is a challenge as these studies are prone to various biases including residual confounding (due to unmeasured or imprecisely measured confounders) and exposure measurement error which can undermine robust causal inference [30, 31].

Mendelian randomization (MR) is an analytical approach that uses randomly assigned (hence unconfounded and pre-disease) germline genetic variants as instruments (i.e., proxies for the risk factor of interest) to examine the causal effects of risk factors on health outcomes [32, 33]. MR is a form of instrumental variable (IV) analysis that allows for unbiased causal effects to be estimated if three assumptions are met: 1) the instrument (e.g., a single germline genetic variant or a multi-allelic score) is robustly associated with the exposure of interest; 2) the instrument is not associated with any confounding factor(s) that would otherwise distort the association between the exposure and outcome; and 3) there is no pathway through which an instrument influences an outcome except through the exposure (known as the “exclusion restriction criterion”). The random allocation of genetic variants at conception and the independent assortment of parental alleles at meiosis means that, at a population level, analyses using genetic variants as instruments for a risk factor of interest should not be confounded by environmental and lifestyle factors that typically distort observational studies.

The availability of germline genetic variants (SNPs – single nucleotide polymorphisms) robustly associated with serum calcium and prostate cancer in separate and independent genome-wide association studies (GWAS) [34, 35] can permit examination of the causal effect of increased serum calcium on prostate cancer risk using a “two-sample Mendelian randomization” framework [36]. Such an approach provides an efficient and statistically robust method of appraising causal relationships between traits, bypassing the need to have access to complete phenotypic and genotypic data on all participants in one sample.

Given uncertainty surrounding the role of serum calcium in prostate cancer aetiology and progression, we used data from: i) a GWAS of serum calcium in up to 61,079 individuals of European descent; and ii) a GWAS of prostate cancer in men of European descent (N=72,729). These samples were used to perform a two-sample Mendelian randomization analysis to examine the causal effect of elevated serum calcium with risk of overall and advanced prostate cancer

## Methods

### Prostate cancer population

We obtained summary genome-wide association study (GWAS) statistics from analyses on 44,825 men with prostate cancer and 27,904 control men of European descent from 108 studies in the Prostate Cancer Association Group to Investigate Cancer Associated Alterations in the Genome (PRACTICAL) consortium [35]. Summary statistics were also obtained from analyses on 6,263 men with advanced prostate cancer (defined as Gleason score ≥8, prostate-specific antigen >100 ng/mL, metastatic disease (M1), or death from prostate cancer) and 27,235 controls. All studies in PRACTICAL have the relevant Institutional Review Board approval from each country, in accordance with the Declaration of Helsinki. Genotype data were obtained by either direct genotyping using an Illumina Custom Infinium array (OncoArray) consisting of approximately 530,000 SNPs [37] or by imputation with reference to the 1000 Genomes Project Phase Three dataset [38]. All SNPs with a poor imputation quality (r^2^<0.30), a minor allele frequency of <1%, a call rate of <98%, or evidence of violation of Hardy-Weinberg equilibrium (*P*<10^−7^ in controls or *P*<10^−^ 12 in cases) were removed. Analyses were performed across individual studies in PRACTICAL using logistic regression in models that were adjusted for the first seven principal components of ancestry (to control for population stratification) and study relevant covariates. Results were meta-analyzed across the PRACTICAL studies using an inverse-variance fixed-effects approach to give an overall effect-estimate.

### Calcium-associated SNP selection

SNPs to proxy for serum calcium were obtained from a GWAS meta-analysis of 39,400 individuals of European descent from 17 population-based cohorts [34]. Genetic instruments were constructed by obtaining SNPs shown to robustly (*P*<1×10^−7^) and independently to associate (*r*^2^<0.01) with serum calcium levels that were replicated (one-sided *P*<0.05) in an independent meta-analysis of up to 21,679 individuals of European descent. In total, 7 SNPs located in or near *CASR* (rs1801725), *DGKD* (rs1550532), *GCKR (*rs780094), *GATA3* (rs10491003), *CARS* (rs7481584), *DGKH* (rs7336933), and *CYP24A1*(rs1570669) were independently replicated. Summary data on rs1801725 were not available in the PRACTICAL OncoArray analysis so we used a proxy SNP located in *CASR* (rs17251221) in high linkage disequilibrium with rs1801725 (*r*^2^=0.85), using the 1000 Genomes Project CEU database as a reference [39]. As an initial test for horizontal pleiotropy (a single locus influencing multiple phenotypes through independent biological pathways; a violation of the “exclusion restriction criterion”), we examined associations of calcium SNPs with thousands of other traits in a large catalogue of summary genetic association statistics from previously published GWAS (MR-Base; www.mrbase.org) [40]. After applying a Bonferroni correction to account for multiple “look ups” of phenotypic traits with all 7 SNPs examined (*P* < 0.05/x, where x represents the number of phenotypic trait “look ups” performed; 859 to 1060 look-ups performed with corresponding corrected *P*-value thresholds: 5.8 x 10^−5^ to 4.7 x 10^−5^ across 7 SNPs), we identified two SNPs (rs780094, rs1550532) that associated with multiple traits in MR-Base. rs780094 was robustly associated (*P*<4.8×10^−5^) with various measures of lipids, insulin, and anthropometric traits and rs1550532 was robustly associated (*P*<4.8×10^−5^) with inflammatory bowel disease; these traits have all been hypothesized to influence prostate cancer risk [41-44]. Additionally, rs1550532 was strongly associated with levels of multiple “unknown metabolites” from untargeted GWAS of metabolomic studies [45]. Given that these two SNPs could influence prostate cancer risk through biological pathways independent of calcium (i.e. horizontal pleiotropy), we removed them from our genetic instrument. Consequently, our genetic instrument for calcium used five SNPs that we assessed as being exclusively associated with serum calcium (rs17251221, rs10491003, rs7481584, rs7336933, rs1570669).

### Statistical analysis

We generated estimates of the proportion of variance in serum calcium for our genetic instrument (R^2^) and F-statistics to examine the strength of our instruments and to test for weak instrument bias (a reduction in statistical power to reject the null hypothesis when an instrument explains only a small proportion of variance in an exposure), using methods previously described [46]. Power calculations were performed using previously reported methods [47] to determine whether we had sufficient sample size to identify effect sizes in our MR analyses that were of a similar magnitude to those reported in the observational literature.

We first examined the effect of serum calcium on overall and advanced prostate cancer for individual SNPs, using the Wald ratio to generate beta-coefficients, and the delta method approximation of the standard error. SNPs were then combined into a multi-allelic genetic instrument (to increase the variance explained in serum calcium) and the causal effect of this instrument on overall and advanced prostate cancer was examined using a maximum likelihood-based approach [48]. For both individual-SNP and multi-allelic instrument analyses, the effect of serum calcium on prostate cancer was scaled to represent a 0.5 mg/dL increase (~ 1 SD). *I*^2^ statistics were calculated to determine the percentage of heterogeneity across SNPs in causal estimates due to variability beyond chance and Cochran’s Q test was used to test homogeneity across SNPs in causal estimates [49]. Maximum-likelihood estimates were then generated using fixed-effects or random-effects models depending on heterogeneity of causal effect estimates across SNPs in multi-allelic instruments. *P*-values were generated using a t-distribution with N-1 degrees of freedom where N is the number of SNPs utilized in the instrument.

To examine the presence of directional pleiotropy (where the horizontally pleiotropic effect across a genetic instrument do not average to zero) from unmeasured traits we performed two sensitivity analyses: MR-Egger regression [50] and the weighted median estimator approach [51]. MR-Egger relaxes the exclusion restriction criterion and thus can provide unbiased estimates of causal effects even when all IVs in an instrument are invalid through violation of this assumption. This approach performs a weighted generalized linear regression of the SNP-outcome coefficients on the SNP-exposure coefficients with an unconstrained intercept term. Provided that the InSIDE (Instrument Strength Independent of Direct Effect) assumption is met (that no association exists between the strength of gene-exposure associations and the strength of bias due to horizontal pleiotropy) and that measurement error in the genetic instrument is negligible (“No Measurement Error” or NOME assumption) [52], the slope generated from MR-Egger regression can provide an estimate of the causal effect of calcium on prostate cancer that is adjusted for directional pleiotropy and the intercept term can provide a formal statistical test for directional pleiotropy. To test NOME, we generated weighted *I*^*2*^_*GX*_ values for overall and advanced prostate cancer analyses to quantify the expected dilution of MR-Egger estimates due to NOME violations [52]. The weighted median estimator (WME) approach provides an estimate of the weighted median of a distribution in which individual IV causal estimates in an instrument are ordered and weighted by the inverse of their variance. Unlike MR-Egger which can provide an unbiased causal effect even when all IVs are invalid, WME requires that at least 50% of the information in a multi-allelic instrument is coming from SNPs that are valid IVs in order to provide an unbiased estimate of a causal effect in an MR analysis. However, the WME has two advantages over MR-Egger in that it provides improved precision as compared to the latter and does not rely on the InSIDE assumption.

We also performed a leave-one-out permutation analysis to examine whether any of our results were driven by any individual SNP from our multi-allelic instrument.

All statistical analyses were performed using R version 3.3.1.

## Results

Our genetic instrument explained 0.71% of variance in serum calcium levels. The corresponding F-statistic for our instrument (86.2) suggested that our instrument was unlikely to suffer from weak instrument bias [53]. Power calculations suggested that we would have 80% power to detect an OR of at least 1.25 (or, conversely a protective OR of at least 0.80) per 0.5 mg/dL increase in serum calcium on overall prostate cancer risk at an alpha level (false positive) of 5%. For advanced prostate cancer, we had 80% power to detect an OR of at least 1.81 (or a protective OR at least 0.55), which would be of similar magnitude to effect estimates reported in the largest observational study of fatal prostate cancer to date (HR [Hazard Ratio] 1.66 per 0.5 mg/dL increase in serum calcium) [26].

Estimates of causal effects of individual calcium SNPs per 0.5 mg/dL increase in serum calcium on overall and advanced prostate cancer per are presented in Table 1. Individually, there was little evidence that any of the 5 SNPs were causally associated with overall or advanced prostate cancer.

**Table 1.** Descriptive statistics of calcium SNPs and estimates of their causal effects on overall and advanced prostate cancer in PRACTICAL.

| SNP | Chr | Gene(s) | EA | NEA | Overall<br>Prostate cancer<br>OR (95% CI) | *P*<br>value | Advanced<br>Prostate cancer<br>OR (95% CI) | *P*<br>value |
| --- | --- | --- | --- | --- | --- | --- | --- | --- |
| rs17251221 | 3 | *CASR* | G | A | 0.84 (0.65-1.09) | 0.18 | 0.83 (0.51-1.35) | 0.45 |
| rs10491003 | 10 | *GATA3* | T | C | 0.56 (0.28-1.15) | 0.12 | 1.58 (0.42-5.93) | 0.50 |
| rs7481584 | 11 | *CARS* | G | A | 0.72 (0.37-1.40) | 0.33 | 1.21 (0.34-4.23) | 0.77 |
| rs7336933 | 13 | *DGKH;*<br>*KIAA0564* | G | A | 1.36 (0.68-2.70) | 0.39 | 1.60 (0.44-5.80) | 0.48 |
| rs1570669 | 20 | *CYP24A1* | G | A | 0.66 (0.35-1.25) | 0.20 | 1.01 (0.31-3.29) | 0.99 |

Chr: Chromosome, EA: Effect Allele, NEA: Non-Effect Allele, OR: Odds Ratio, 95% CI: 95% Confidence Interval. EA reflects the allele that increases serum calcium levels. OR (95% CI) represents the exponential increase in odds for each 0.5 mg/dL increase in serum calcium

### Overall prostate cancer

In an MR analysis combining the five serum calcium-related SNPs into a multi-allelic genetic instrument, there was weak evidence of a protective effect of serum calcium on prostate cancer risk (OR per 0.5 mg/dL increase in calcium: 0.83, 95% CI: 0.63-1.08; *P*= 0.12) (Table 2). Effect estimates were similar using the weighted median estimator (OR 0.80, 95% CI: 0.58-1.12) and MR-Egger (OR 0.87, 95% CI: 0.46-1.64). The MR-Egger intercept parameter did not suggest evidence of directional pleiotropy (OR 1.00, *P*=0.76).

**Table 2.** Mendelian randomization derived causal effects of a 0.5 mg/dL increase in serum calcium on overall and advanced prostate cancer using a multi-allelic instrument in PRACTICAL.

|  | **Maximum**<br>**likelihood**<br>**estimate**<br>**OR (95% CI)<sup>a</sup>** | **Weighted median**<br>**estimator**<br>**OR (95% CI)** | **MR-Egger**<br>**regression**<br>**OR (95% CI)** | **MR-Egger**<br>**regression**<br>**intercept term**<br>***P*-value** |
| --- | --- | --- | --- | --- |
| **Overall prostate**<br>**cancer (N=72,729)<sup>b</sup>** | 0.83 (0.63-1.08) | 0.80 (0.58-1.12) | 0.87 (0.46-1.64) | 0.76 |
| **Advanced prostate**<br>**cancer (N=33,498)<sup>c</sup>** | 0.98 (0.57-1.70) | 0.92 (0.50-1.66) | 0.72 (0.24-2.15) | 0.42 |

<sup>a</sup> Odds ratio [OR] (95% confidence interval, CI) represents the exponential increase in odds for each 0.5 mg/dL increase in serum calcium. <sup>b</sup> maximum likelihood estimate obtained using a fixed-effects model ( $I^2=0\%$ , $Qp=0.44$ ); <sup>c</sup> fixed-effects model ( $I^2=0\%$ , $Qp=0.80$ ).

### Advanced prostate cancer

MR analyses found little evidence for an effect of serum calcium on advanced prostate cancer risk (0.5 mg/dL calcium increase: OR 0.98, 95% CI: 0.57-1.70; *P*=0.93) (Table 2). Sensitivity analyses to examine directional pleiotropy were consistent with a null effect of serum calcium on advanced prostate cancer.

Calculation of the *I^2^_GX_* statistic suggested little attenuation of our MR-Egger estimates due to measurement error for both overall prostate cancer (*I^2^_GX_*=0.89) and advanced prostate cancer (*I^2^_GX_*=0.91), so adjustment of MR-Egger estimates to account for mild dilution bias was not performed [52]. Leave-one-out permutation analyses for overall and advanced prostate cancer did not find evidence that the effect estimate based on the multi-allelic instrument was being driven by any single serum calcium related SNP (Supplementary Table 1).

## Discussion

Our Mendelian randomization analysis does not support the hypothesis that serum calcium increases the risk of overall or advanced prostate cancer. Indeed, the point estimates were in the opposite direction (though imprecisely estimated) to findings from some observational studies.

Our findings are not consistent with some laboratory studies which have reported a role of calcium in promoting loss of differentiation and increased proliferation of prostate cancer cells [54, 55]. Further, our results are not consistent with a meta-analysis of prospective observational studies that reported dose-response relationships of dietary calcium intake (per 400 mg/day) with risk of prostate cancer (RR 1.05, 95% CI: 1.02-1.09, N=15 studies), though there was moderate heterogeneity in associations across studies (*I*^*2*^ = 49%, *P*- heterogeneity = 0.02) [11].

Prospective studies that have examined the association of serum calcium with incident prostate cancer have generated conflicting findings: three did not find strong evidence for an association (HR for upper vs. lower tertile: 1.31, 95% CI: 0.77-2.20 [23]; OR for upper vs. lower quartile: 1.04, 95% CI: 0.78–1.39 [28]; HR per quartile increase: 0.99, 95% CI: 0.94– 1.03 [29]), whereas one reported a weak inverse association between calcium and prostate cancer (HR per SD increase: 0.97, 95% CI: 0.85-1.00) [24]. Likewise, some studies that have examined an association between serum calcium and fatal prostate cancer have reported positive risk relationships (HR for upper vs. lower tertile: 2.07, 95% CI: 1.06-4.04 [27]; HR for upper vs. lower tertile: 2.68, 95% CI: 1.02-6.99 [23]; HR per 0.1 mmol/L increase: 1.50, 95% CI: 1.04–2.17 [26]) whereas others have not found strong evidence of an association (HR per 1-SD increase: 1.00, 95% CI: 0.92–1.09 [24]; HR for upper vs. lower quartile: 0.75, 95% CI: 0.49–1.15 [25]). It is plausible that discordance between previously reported observational findings and our MR analysis may reflect residual confounding in the former (e.g., through other dietary, lifestyle, or molecular factors). Nevertheless, the weak evidence that we found for a potential protective effect of serum calcium on overall prostate cancer is consistent with a meta-analysis of four randomized controlled trials that reported that daily calcium supplementation (≥ 500 mg/day) reduced prostate cancer risk (RR [Risk Ratio] 0.54, 95 % CI: 0.30-0.96, *P*= 0.03), though this analysis was only based on 48 men with prostate cancer (3297 and 3248 in the intervention and control groups, respectively) [56].

Strengths of our analysis include the use of a Mendelian randomization approach to appraise the relationship of serum calcium with prostate cancer risk which should help to minimize or avoid confounding through lifestyle or environmental factors that may have biased findings from previous observational analyses. Further, given the time required for nutritional biomarkers to influence carcinogenesis [57] and the considerable latency period of prostate cancer [58], the use of germline genetic variation as an instrument should allow for sufficient time to confer an effect on prostate cancer. This is because MR will estimate the effect of life-long exposure to elevated serum calcium on prostate cancer risk. MR will also offer an additional strength over prospective studies of dietary or serum calcium which can suffer from substantial (albeit, likely non-differential) measurement error: measurement error in genetic studies is often low as modern genotyping technologies provide relatively precise measurement of genetic variants [59]. The use of a two-sample MR approach allowed us to utilize summary effect estimates from two large GWAS and thus increase statistical power in our analyses. Additionally, though the F-statistic generated for our instrument suggested that weak instrument bias was unlikely, in a two-sample MR setting, weak instrument bias if present would be expected to bias associations toward the null, providing a conservative effect estimate. This is in contrast to a one-sample MR analysis in which weak instrument bias will tend to bias effect estimates toward the confounded observational study estimate [36]. Lastly, by obtaining summary effect estimates for both exposure and outcome datasets from GWAS that were restricted to individuals of European descent and adjusted for principal components of ancestry, we reduced (though did not eliminate) the possibility of confounding through population stratification in our MR analyses (though this may limit generalizability of our findings to other ethnicities).

There are limitations to our analysis. First, given the composite characterization of advanced prostate cancer in the summary GWAS data that we obtained (Gleason ≥8, prostate-specific antigen >100 ng/mL, metastatic disease (M1), or death from prostate cancer), it is difficult to directly compare our findings with those from prospective studies that examined associations between calcium and fatal prostate cancer. Second, though our MR analysis for advanced prostate cancer was sufficiently powered to detect effect sizes compatible with those reported in the observational literature, it was not powered to detect effect sizes of a more modest magnitude. Further identification of independent genetic variants that influence serum calcium (increasing instrument strength further by explaining a larger proportion of the variance in serum calcium) in addition to larger GWAS of advanced prostate cancer will help to improve statistical power for future analyses. A final limitation of our analysis was that we were unable to examine possible non-linear effects of serum calcium on prostate cancer using summarized genetic data, which have been proposed previously [8].

Given that our findings raise the possibility that serum calcium may be protective against prostate cancer, there is a need to follow-up these results in large and independent datasets. Further identification of additional independent genetic variants robustly associated with serum calcium will help to improve precision of future analyses.

In conclusion, our Mendelian randomization analysis does not support the hypothesis that serum calcium increases the risk of overall or advanced prostate cancer.

## Acknowledgments

The authors’ responsibilities were as follows—SJL and CB: conceived the study; JY, KB, RL, CB, and SL: planned the analyses; JY: conducted data analysis; JY, KB, RL: prepared the manuscript; CB, the PRACTICAL consortium, GDS, RMM, SL: critically revised the manuscript; and all authors: read and approved the final manuscript. None of the authors had any conflicts of interest to declare.

